# Multi-hierarchical Profiling the Structure-Activity Relationships of Engineered Nanomaterials at Nano-Bio Interfaces

**DOI:** 10.1101/273268

**Authors:** Xiaoming Cai, Jun Dong, Jing Liu, Huizhen Zheng, Chitrada Kaweeteerawat, Fangjun Wang, Zhaoxia Ji, Ruibin Li

## Abstract

Increasingly raised concerns (nanotoxicity, clinical translation, *etc*) on nanotechnology require breakthroughs in structure-activity relationship (SAR) analyses of engineered nanomaterials (ENMs) at nano-bio interfaces. However, current nano-SAR assessments failed to disclosure sufficient information to understand ENM-induced bio-effects. Here we developed a multi-hierarchical nano-SAR assessment for a representative ENM, Fe_2_O_3_ by systematically examining cellular metabolite and protein changes. This nano-SAR profile allows visualizing the contributions of 7 basal properties of Fe_2_O_3_ to their diverse bio-effects. For instance, while surface reactivity is responsible for Fe_2_O_3_-induced cell migration, the inflammatory effects of Fe_2_O_3_ nanorods and nanoplates are determined by their aspect ratio and surface reactivity, respectively. We further discovered the detailed mechanisms, including NLRP3 inflammasome pathway and monocyte chemoattractant protein-1 involved signaling. Both effects were further validated in animal lungs. Our findings provide substantial new insights at nano-bio interfaces, which may facilitate the tailored design of ENMs to endow them with desired bio-effects.

---

The physicochemical properties of engineered nanomaterials (ENMs) have been demonstrated to play a decisive role in nano-bio interactions^1^. Given the rapidly increasing number of ENMs as well as their diverse physicochemical properties including size, shape, surface area, surface reactivity, mechanical strength, *etc.*^2^, the *in vitro* structure-activity relationship (SAR) studies on ENMs have significantly promoted the development of nanobiotechnology^3-5^. In general, nano-SAR analyses have enabled the determination of the key physicochemical properties of ENMs that are responsible for evoking a target bio-effect in the organism^1, 6^, allowed bio-hazard ranking of various new ENMs^7^, and facilitated the engineering design of biocompatible materials by tailored functionalization ^8^. However, current nano-SAR analyses only focus on the influence of a single property (size, charge, or surface charge, *etc.*) of ENMs to individual bio-effects (*e.g.* apoptosis, necrosis, autophagy or inflammation, *etc*.)^2^. Considering some increasingly raised bottleneck problems in nanotechnology, e.g. various ENM-induced nanotoxicities^3, 4^, and severe clinical translation barriers in nanomedicine^10^, there is a demand for tiered views of nano-SARs.

System biology is a new theme in biological science, aiming at system-level understanding of biological organisms. Several omics-based technologies including genomics, proteomics, metabolomics, *etc*., have been developed for systematic analysis of biomolecules (genes, proteins, metabolites, *etc*.) expressed in cells or tissues^10^. Recently, some progresses have been made using omics to investigate protein corona on ENM surfaces^11^, examine ENM-induced cell signaling changes^12, 13^, define the routes of ENM trafficking^14^, decipher cytotoxicity mechanisms^15^, *etc*. However, so far, no attempts have been made for nano-SAR assessments^16^. Since proteins and metabolites are the executors or end products of signaling pathways and multi-omics analyses offer a better view of the global biological changes^17^, we hypothesized that multi-hierarchical nano-SAR assessments could be achieved *via* coupling of proteomics and metabolomics analyses.

In this study, we engineered a series of iron oxide nanoparticles to assess their SAR because they are widely used in constructions^18^, pigments^19^, biomedicine^20,21^, *etc*, and their global production had reached to 1.83 billion in 2015. We selected Fe_2_O_3_ nanorods and nanoplates here based on our previous experience that various nanorods like CeO_2_, AlOOH and lanthanide materials, or nanoplates (e.g. Ag nanoplates) were demonstrated to be more reactive than other shapes^22-24^. The metabolomics and proteomics changes induced by Fe_2_O_3_ particles were examined in THP-1 cells, a macrophage-like cell line, which are the first port of entry for the ENMs exposed to mammalian systems^7, 25^. A multi-hierarchical nano-SAR profile was established by integration of the physicochemical properties of Fe_2_O_3_ particles, biological effects and their correlation coefficients. The identified nano-SARs were selectively validated by deciphering the detailed mechanisms *in vitro* and *in vivo*.

## Results

### Preparation and characterization of Fe_2_O_3_ nanoparticle library

To explore the SAR of Fe_2_O_3_, we synthesized a series of Fe_2_O_3_ NPs with different morphologies and sizes, including 4 hexagonal nanoplates (P1∼P4) with controlled diameters and thicknesses, and 4 nanorods (R1∼R4) with systematically tuned lengths and diameters. Transmission electron microscopy (TEM) was used to determine the size and morphology of all Fe_2_O_3_ particles. Figure 1A shows that the diameters of Fe_2_O_3_ nanoplates range from 45 to 173 nm and their thicknesses are 16∼44 nm, while the lengths and diameters of nanorods are 88∼320 and 20∼53 nm, respectively. We further calculated the ratios of diameter to thickness for the nanoplates, length to diameter for nanorods, respectively, and denoted them as aspect ratios (ARs). The ARs of Fe_2_O_3_ nanoplates and nanorods are 1.0∼10.8 and 1.7∼8.0, respectively. The surface areas were 16∼27 m^2^/g, determined by Brunauer-Egmmett-Teller (BET) method (Table 1).

**Table 1.** Quantitative Characterization of Fe_2_O_3_ nanoparticles

| Properties | Detection methods | Quantitative Characterization |  |  |  |  |  |  |  |
| --- | --- | --- | --- | --- | --- | --- | --- | --- | --- |
|  |  | Plates |  |  |  | Rods |  |  |  |
|  |  | P1 | P2 | P3 | P4 | R1 | R2 | R3 | R4 |
| Length (nm) | TEM | NA | NA | NA | NA | 88 ± 8 | 181 ± 11 | 116 ± 12 | 322 ± 26 |
| Diameter (nm) | TEM | 45 ± 3 | 84 ± 5 | 122 ± 7 | 173 ± 6 | NA | NA | NA | NA |
| Thickness (nm) | TEM | 44 ± 7 | 23 ± 3 | 18 ± 2 | 16 ± 3 | 53 ± 8 | 38 ± 5 | 20 ± 4 | 40 ± 7 |
| Aspect ratio | Cal.* | 1.0 ± 0.1 | 3.7 ± 0.2 | 6.8 ± 0.1 | 10.8 ± 0.3 | 1.7 ± 0.2 | 4.5 ± 0.4 | 5.8 ± 0.5 | 8.0 ± 0.7 |
| Hydrodynamic size (nm) | DLS | 175 ± 10 | 246 ± 7 | 366 ± 16 | 378 ± 8 | 422 ± 12 | 463 ± 8 | 578 ± 8 | 536 ± 4 |
| Zeta potential (nm) | ZPA | -7.9 ± 0.8 | -3.6 ± 0.6 | -5.1 ± 0.4 | -3.7 ± 0.7 | -6.5 ± 0.7 | -10.7 ± 1.6 | -8.5 ± 0.6 | -5.2 ± 2.1 |
| Surface area (m<sup>2</sup>/g) | BET | 22 ± 1.6 | 17.9 ± 1.0 | 17.4 ± 0.6 | 16.8 ± 0.4 | 18.3 ± 1.5 | 20.8 ± 1.8 | 26.9 ± 2.3 | 21.3 ± 1.8 |
| Surface activity (a.u.) | DCF | 1.25 ± 0.08 | 2.59 ± 0.11 | 3.23 ± 0.15 | 1.46 ± 0.08 | 0.59 ± 0.04 | 0.35 ± 0.03 | 0.63 ± 0.07 | 0.47 ± 0.03 |

\*Calculation (Cal.): Aspect ratio = $\frac{Length\ (or\ Diameter)}{Thickness}$

**Figure 1.**
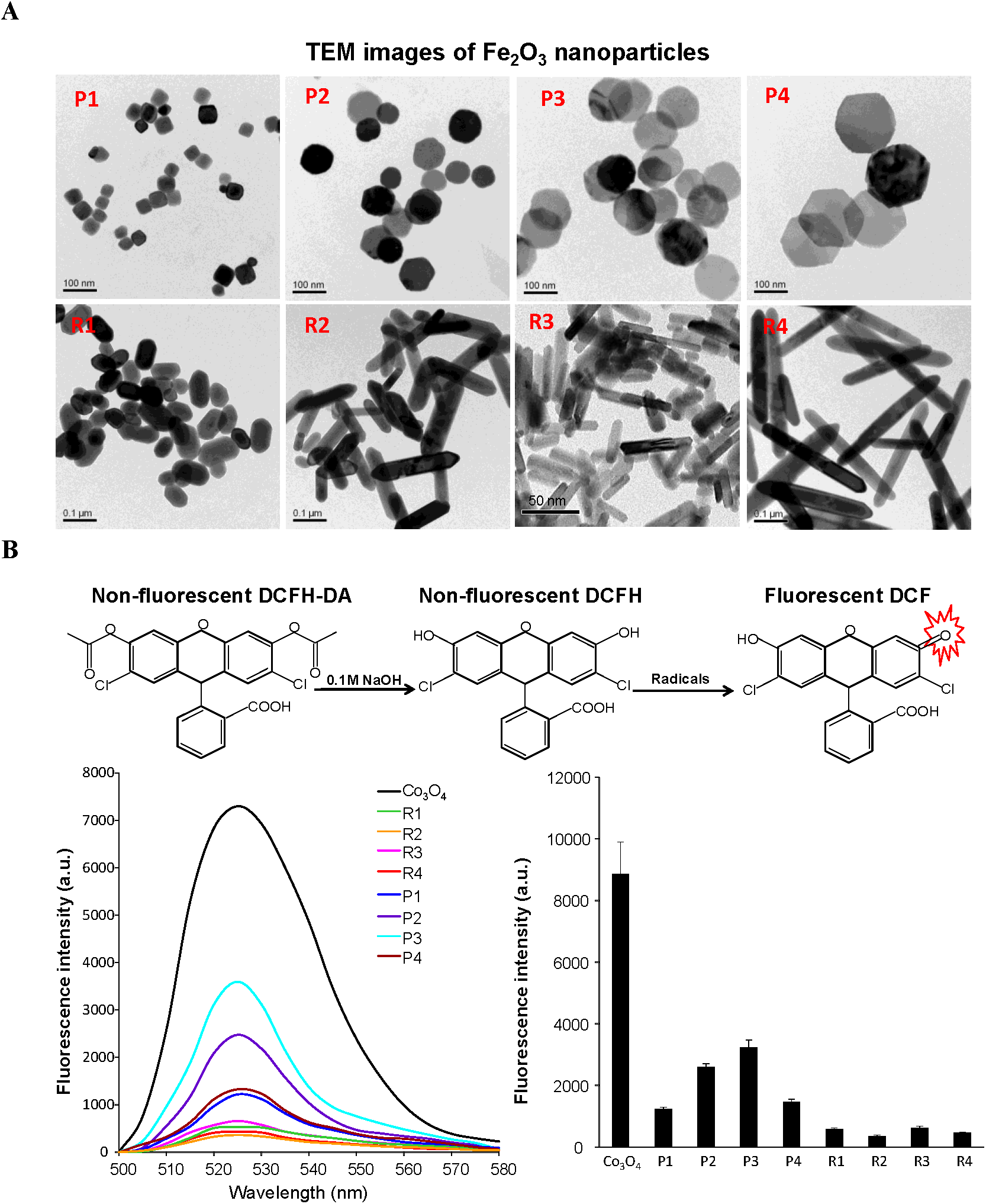
Characterization of Fe_2_O_3_ nanoparticles by TEM and DCF assay. A) TEM images and B) surface reactivity of Fe_2_O_3_ nanoparticles. TEM samples were prepared by placing a drop of the particle suspensions (50 μg/mL in DI H_2_O) on the grids. To assess the surface reactivity of Fe_2_O_3_ samples, 95 μL aliquots of 25 ng/mL DCF were added into each well of a 96-multiwell black-bottom plate and mixed with 5 μL of nanoparticle suspensions at 5 mg/mL, followed by 2 h incubation. A SpectraMax M5 microplate reader was used to record the fluorescence emission spectra of DCF agent at an excitation wavelength of 490 nm.

X-ray diffraction analysis (XRD) was performed to determine the crystal structure of Fe_2_O_3_. Figure S1 shows the XRD patterns for selected nanoplates and nanorods, P2 and R2. All diffraction peaks could be indexed to rhombohedral α-Fe_2_O_3_ phase (JCPDS no. 33-0664) and no other impurity peaks were detected.

The hydrodynamic sizes in RPMI medium were assessed by dynamic light scattering (DLS), showing ranges from 170 to 380 nm for plates and 420-540 nm for rods (Table 1). The surface charges of Fe_2_O_3_ particles were determined by a Zeta potential analyzer (ZPA), and showed very similar Zeta potential values of 0 to −10 mV in cell culture media, which reflects the formation of protein corona on particle surface.

We used the 2’,7’-dichlorofluorescein (DCF) assay to investigate the surface reactivity of Fe_2_O_3_ nanoparticles. The DCF assay is based on a mechanism that nonfluorescent 2’,7’-dichlorodihydrofluorescein (H_2_DCF) could be converted to the highly fluorescent DCF by oxidation. This assay has been widely used to access the radicals or abiotic reactive oxygen species (ROS) generation on nanoparticle surface^6, 8^. Co_3_O_4_ nanoparticles have been demonstrated to exhibit high surface reactivity in DCF assay and were used as a positive control^26^. As shown in Figure 1B, Fe_2_O_3_ nanoplates are more reactive than nanorods, and P3 exhibits the highest surface oxidative capability. These differences in surface reactivity may results from their crystal facets. XRD analysis shows that (104) is the dominant facet in Fe_2_O_3_ nanoplates as compared to (110) as the strongest peak for the nanorods (Figure S1). This is consistent with several earlier studies showing that (104) as the dominant facet of the α-Fe_2_O_3_ nanoplates is highly catalytically active^27^. Therefore, the high surface reactivity of nanoplates observed here can be attributed to the exposure of more active (104) facets.

### Using metabolomics to explore Fe_2_O_3_ induced metabolite changes

Metabolites are the end products of diverse intracellular processes, so the changes of cell metabolome can reflect cell responses to stimuli. We performed a global metabolomics study to explore the metabolic changes induced by Fe_2_O_3_ nanoparticles in THP-1 cells, a myeloid cell line that is often used as an *in vitro* model for studying the effects of engineered nanoparticles on immune cells^28^. As described in the experimental section, the metabolites in THP-1 cells after Fe_2_O_3_ treatment were extracted and subjected to C18 reversed-phase column for nontargeted liquid chromatography-mass spectrometry (LC-MS) analysis on a high-resolution Triple Time of Flight (TOF) mass spectrometry in both positive (ESI+) and negative (ESI-) ionization modes. By use of XCMS software, 8001 and 3479 metabolite features were obtained from the LC-MS data collected in ESI+ and ESI-mode, respectively. One-way analysis of variance (ANOVA) was used to screen metabolite differences associated with Fe_2_O_3_ treatment. The significance of each feature was determined by its p-value and false discovery rate (FDR) truncated at 0.01 and 0.05, respectively. As a result, 1674 and 1180 discriminating features were detected in the data of ESI+ and ESI-mode, respectively. Figure 2A shows a heat map of the significant features. Compared to the control, Fe_2_O_3_-treated samples show increases in most of the detected features. R4 induces the most significant metabolic changes in THP-1 cells, R1, R2, R3 and P3 show moderate effects, while P1, P2 and P4 are relatively bio-inert.

**Figure 2.**
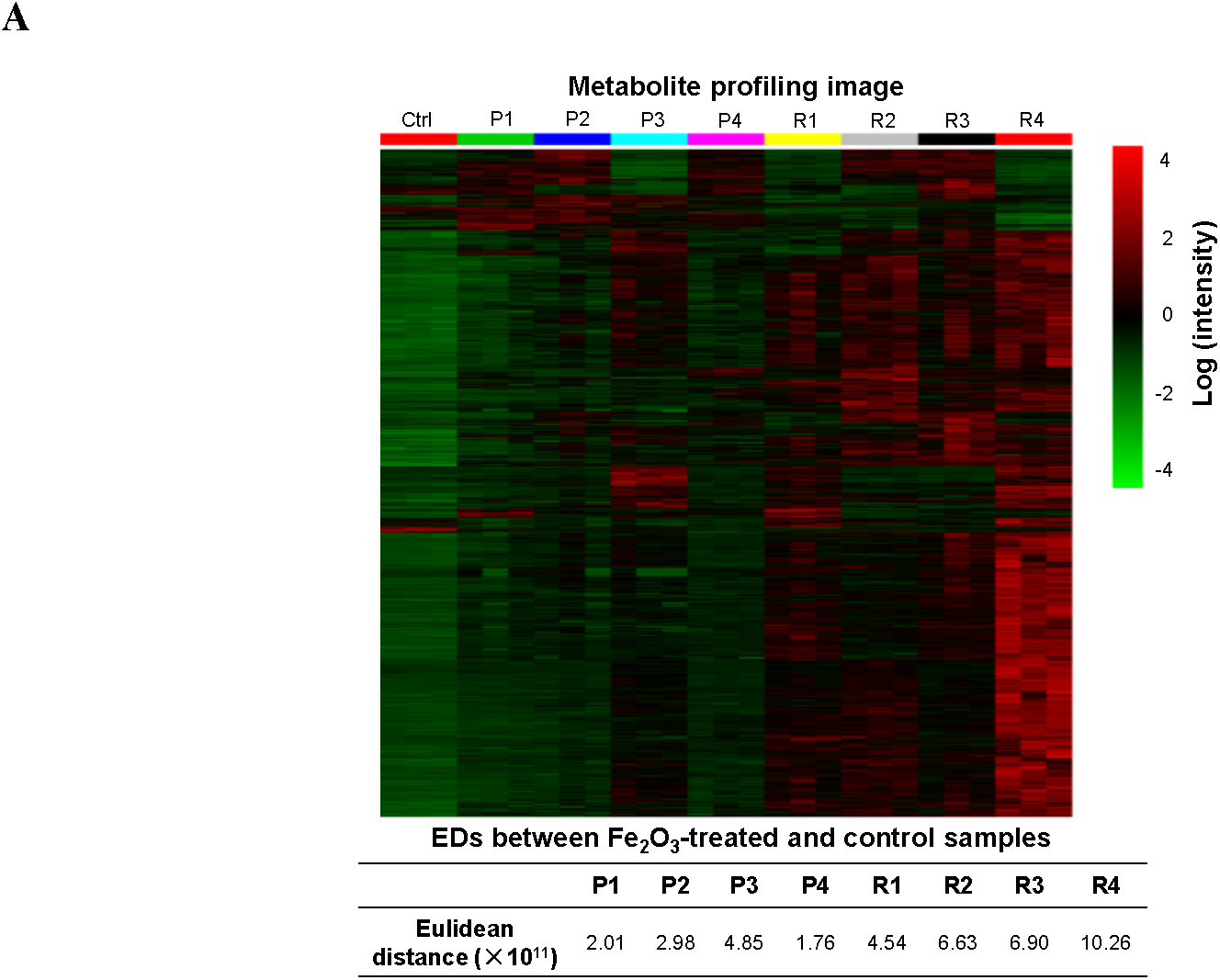
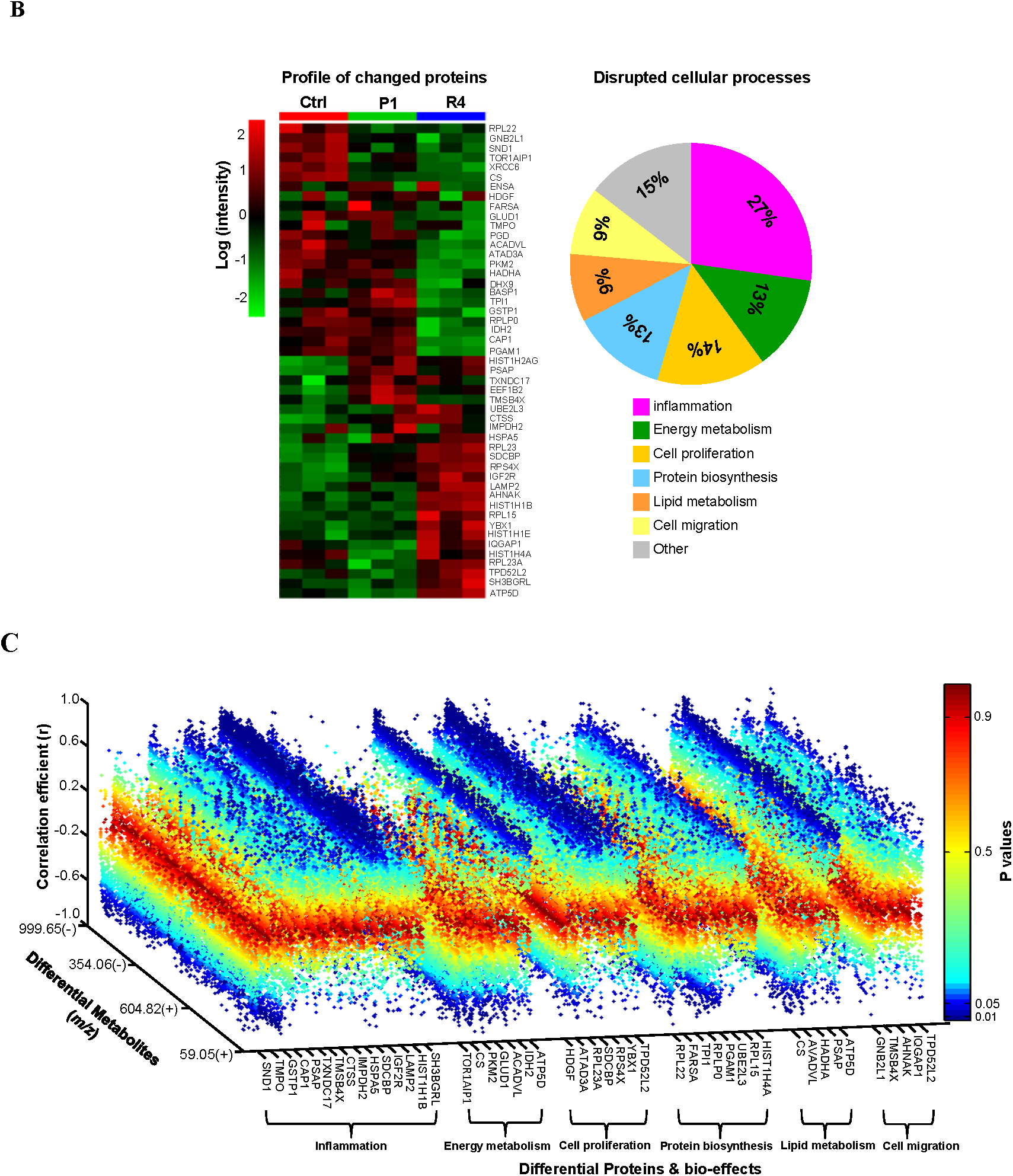
Fe_2_O_3_ induced metabolomics and proteomics changes and their relationships. **A)** Metabolite profile of THP-1 cells exposed to Fe_2_O_3_ library as well as their Euclidean distances (EDs). After 24 h treatment, the cell samples (n= 3 independent experiments) were collected to extract proteins and metabolites. For metabolomics analysis, the log-transformed normalized peak intensities of metabolites in all the cell samples were expressed using red, black or green colors in a heat map. EDs between control and nanoparticle-treated samples were calculated to quantitatively evaluate the global metabolite profile changes induced by Fe_2_O_3_. **B)** Fe_2_O_3_ nanoparticle induced protein expression changes and bio-effects. For proteomics analysis, a heat map was plotted in a similar way as metabolites, showing 49 significantly changed proteins in proteomics analysis (left panel). The pathways or bio-effects related with these proteins were determined by KEGG and UniprotKB database (right panel). The percentages of the differential proteins involved in each specific bio-effects of Fe_2_O_3_ were shown in the pie chart. **C)** Regression analysis between differential proteins and metabolites. The regression analysis of the differential proteins and metabolites was performed in Matlab R2009b. The correlation is considered as statistically significant with correlation coefficient (r) ≤-0.8 or r≥0.8, and p ≤ 0.01.

We further performed a hierarchical clustering analysis of metabolites to calculate the Euclidean distances (EDs) between control sample and nanoparticle-treated samples^29^. This parameter was used to quantitatively describe the global metabolite profile changes by Fe_2_O_3_, and a longer distance usually means more disruptions to the homeostasis of cellular metabolism, which could be considered as a bio-activity index of stimuli at systemic levels^30^. As shown in Figure 2A, the ED ranking of different Fe_2_O_3_ particles is R4> R3> R2> P3> R1> P2> P1> P4, which is consistent with the observation on metabolite heat map. Linear regression analysis was used to investigate the relationships between EDs and the physical chemical properties of Fe_2_O_3_ nanoparticles (Figure S2). According to the r^2^ values of regression models, surface reactivity and aspect ratio are the dominant physicochemical properties for the global metabolite changes in THP-1 cells, and account for 88.98% and 98.02% of ED variations in nanoplates and nanorods, respectively.

### Discovery of the bio-effects of Fe_2_O_3_ particles by proteomics

Proteins, as a major executor of signaling pathways in biological organisms are involved in many cellular effects. The bio-effects of Fe_2_O_3_ nanoparticles in cells could be determined by identification of the proteome changes. Since the metabolomics profile suggests that R4 and P1 are the most bio-active and bio-inert materials, respectively, they were selected and exposed to THP-1 cells for proteomics analysis. The protein expression was analyzed by a nanoscale liquid chromatography coupled to tandem mass spectrometry (nano LC-MS/MS) as described in the experimental section, and 785 proteins were identified for statistical analysis.

To discover the biomarkers related to the bio-effects of Fe_2_O_3_, ANOVA analysis was performed. As a result, 49 identified proteins with p values< 0.01 and FDRs< 0.05 were considered to be significantly changed after Fe_2_O_3_ treatment. The data were further integrated into a heat map to visualize the expression levels of these proteins in control, P1 and R4 samples (Figure 2B). While R4 induced significant proteome changes in THP-1 cells including 25 up-regulated and 24 down-regulated proteins, P1 had negligible effects. We used KEGG and UniprotKB database to investigate the impacts of the 49 proteins to cell pathways and functions. These proteins were found to mainly participate in 6 biological processes, including inflammation, cell proliferation, energy metabolism, lipid metabolism, protein biosynthesis and cell migration (Figure 2B). Regression analysis was used to explore the relationships between the changed proteins and the metabolites. As shown in Figure 2C, correlations between the bio-effects and metabolite changes could be plotted successfully, which provides an opportunity to explore the relationships between the physicochemical properties of Fe_2_O_3_ nanoparticles and their bio-effects.

### Profiling the multi-hierarchical nano-SAR of Fe_2_O_3_ nanoparticles by a 3D heatmap

A 3D heatmap was plotted to quantitatively describe the influences of physicochemical properties to the bio-effects of Fe_2_O_3_ nanoparticles by regression analysis among the properties of nanoparticles, their metabolite changes and bio-effects (Figure 3). While the zeta potential of Fe_2_O_3_ nanoplates has some effects on cell proliferations with coefficient r^2^ value at 0.64, surface reactivity is the dominant property that impacts other 5 bio-effects as well as global cellular changes. For Fe_2_O_3_ nanorods, surface reactivity is responsible for the disruption of cell migration (r^2^=0.88) and protein biosynthesis (r^2^=0.99); particle length significantly affects the energy and lipid metabolism processes; AR plays a major role in inflammation and cell proliferation. These results suggest that there is one dominant property that best correlates with a specific bio-effect. This is the first time that we determined the contributions of 7 basal physicochemical properties of ENMs to their diverse bio-effects by plotting a nano-SAR profile at multi-hierarchical levels.

**Figure 3.**
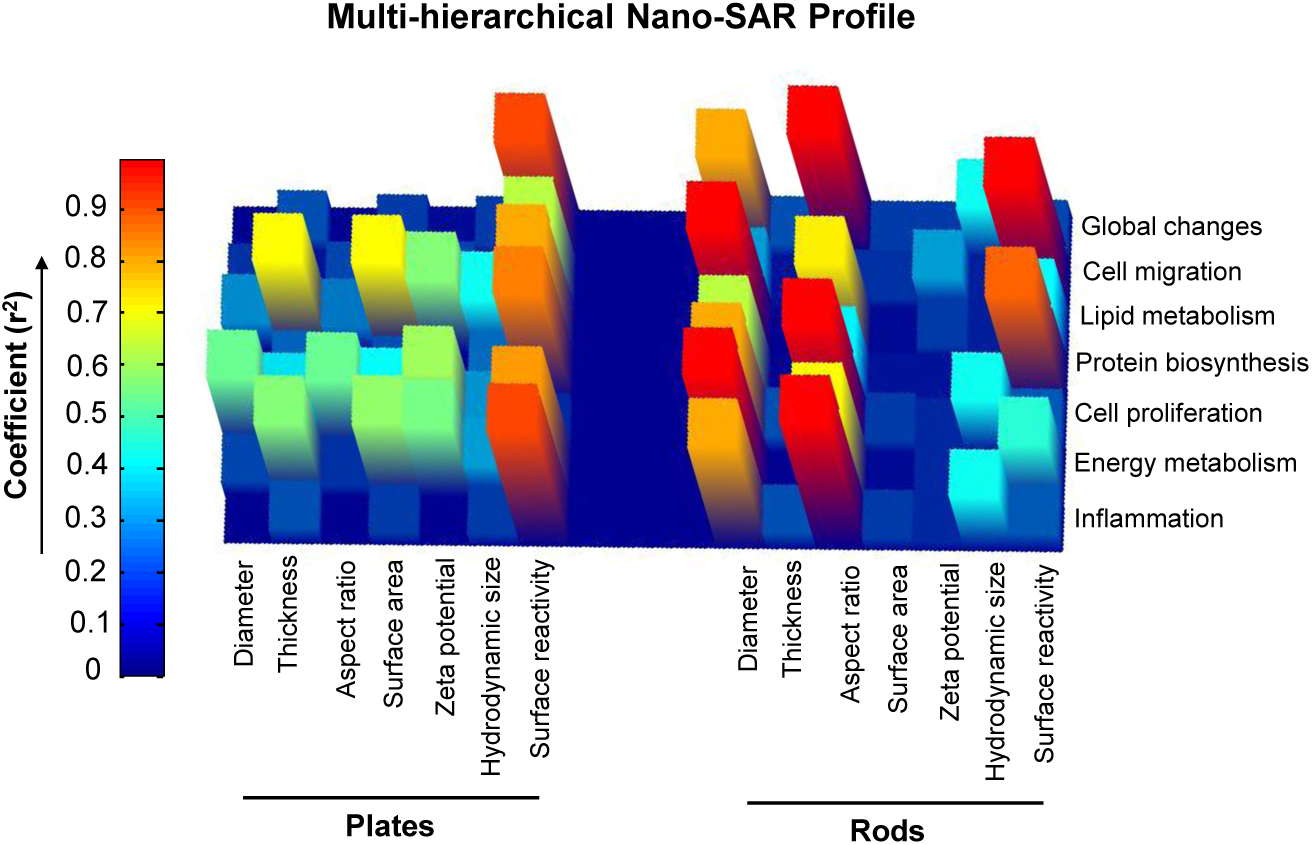
Multi-hierarchical profiling the nano-SAR of Fe_2_O_3_ particles. The relationships between the 7 physicochemical properties of Fe_2_O_3_ nanoparticles and their bio-effects could be visualized by the 3D heatmap, which is established by regression analyses among metabolites, differential proteins and Fe_2_O_3_ properties.

### Exploring the inflammatory effects of Fe_2_O_3_ nanoparticles in THP-1 cells by deciphering the detailed mechanism

The nano-SAR profile indicates that Fe_2_O_3_ nanoparticles may induce significant inflammatory effects in THP-1 cells. Among the changed proteins, 15 of them are involved in inflammation pathways, *e.g.*, proactivator polypeptide (PSAP), cathepsin S (CTSS), and cation-independent mannose-6-phosphate receptor (IGF2R). These proteins have effects on phagocytosis and lysosomal dysfunction^31-33^, implying a lysosome-involved mechanism. We further investigated this by detecting the pro-inflammatory cytokine release in THP-1 cells. Monosodium urate (MSU) was used as a positive control to evoke inflammatory response. Although Fe_2_O_3_ nanoparticles have little effect on cytokine production in THP-1 cells exposed to 0-100 μg/mL particles for 24 h (Figure S3A), all Fe_2_O_3_-treated cells exhibit significant IL-1β and TNF-α increase in dose-dependent manners at 48 h (Figure 4A and S3B). However, all these particles show little effects in cell viability (Figure S4). At the 100 μg/mL exposure dose, R4 exhibits the highest inflammatory cytokine production, P3, R2 and R3 have moderate effects, while P1, P2, P4 and R1 induce a small amount of cytokine release. This trend can’t be explained by the cellular uptake levels of Fe_2_O_3_ nanoparticles. Although the nanorods in general have relatively higher cellular uptake than the nanoplates, there’s no difference among nanorods (or nanoplates) (Figure S5). Consistent with the nano-SAR profile, cytokine productions by rods and plates have good correlation with their aspect ratios and surface reactivity, respectively.

**Figure 4.**
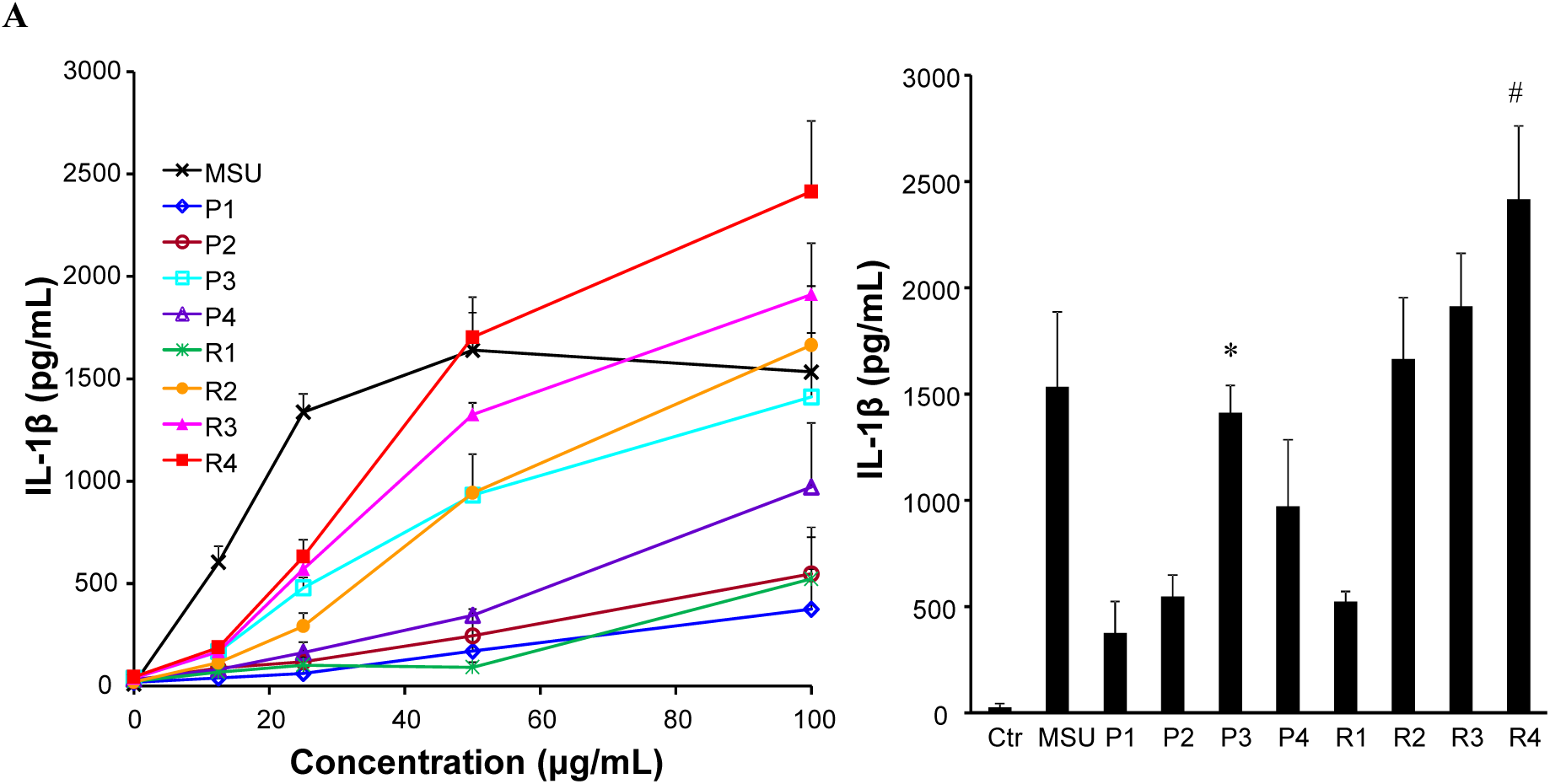
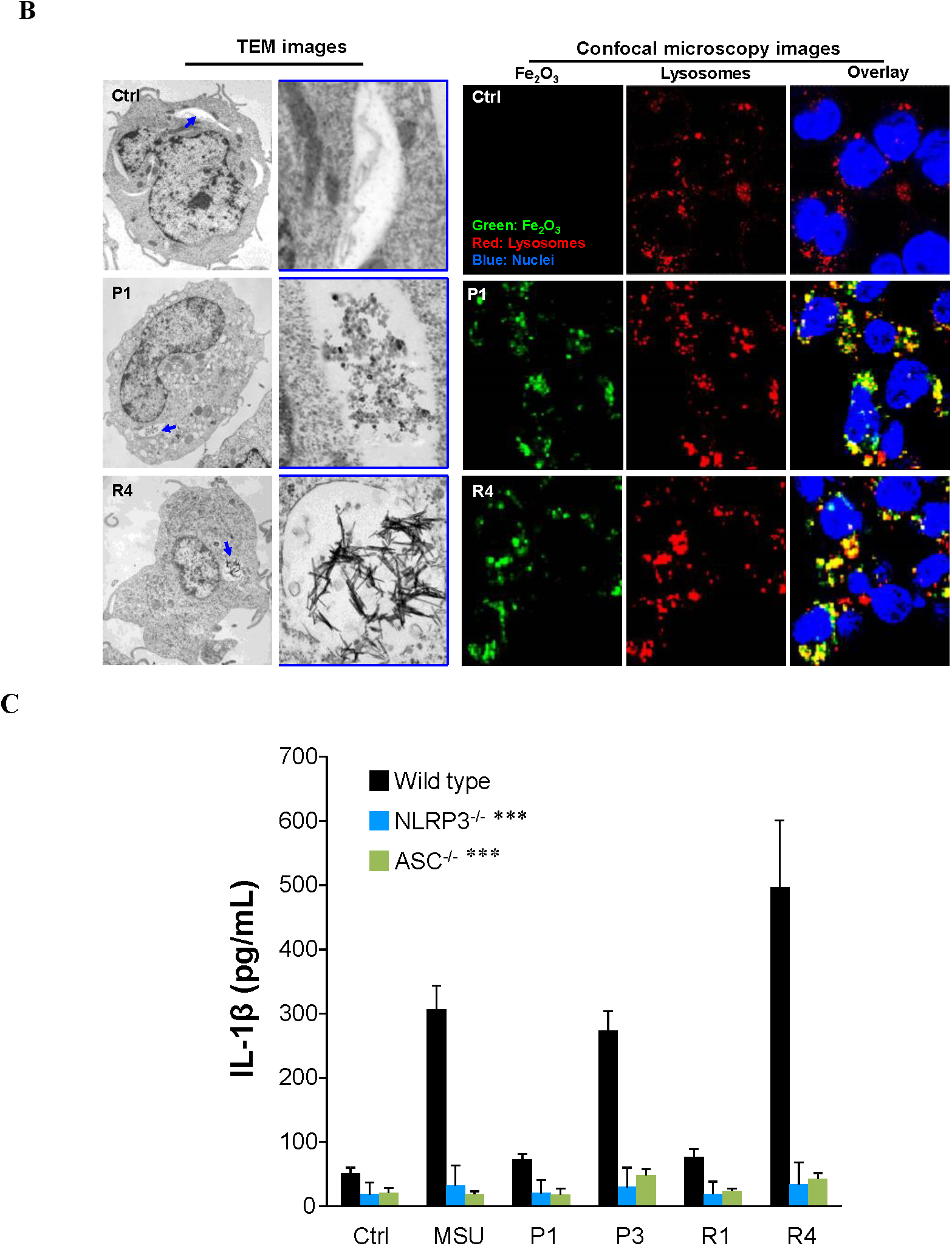
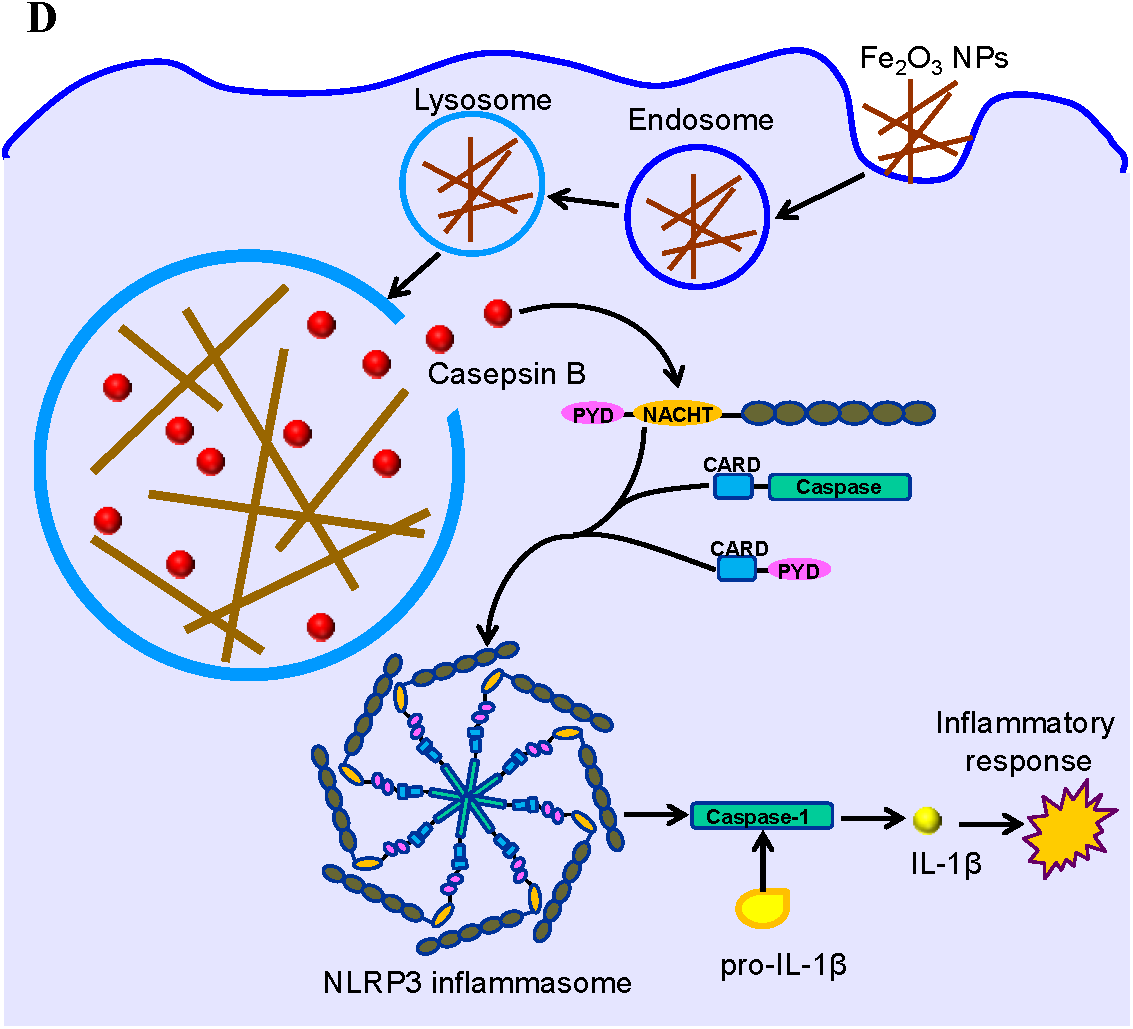
Determination of the inflammatory pathway of Fe_2_O_3_ in THP-1 cells. **A)** IL-1β production in THP-1 cells exposed to 0-100 μg/mL Fe_2_O_3_ nanoparticles for 48 h. *p< 0.05 compared to P1,P2 and P4, #p< 0.05 compared to other particles (two-tailed Student’s t-test). After 48 h incubation of THP-1 cells with Fe_2_O_3_ particles, the supernatants were collected to quantify cytokine productions by ELISA. **B)** TEM and confocal microscopy imaging of internalized Fe_2_O_3_ nanoparticles. THP-1 cells exposed to pristine or FITC-labeled Fe_2_O_3_ nanoparticles were washed, fixed and stained for TEM or confocal microscopy imaging. Hoechst 33342 dye (blue) and Alexa fluor 594 labeld antibodies (red) were used to identify nuclei and lysosomes, respectively. **C)** comparison of IL-1β production in Wild-type, NALP3^-/-^, and ASC^-/-^ THP-1 cells exposed to Fe_2_O_3_ nanoparticles, ***p< 0.001 compared to particle-treated Wild-type cells (two-tailed Student’s t-test). Data in the line and bar graphs are shown as mean ± s.d. from 4 independent replicates. **D)** schematic to explain the inflammatory mechanism.

In order to understand the detailed mechanisms involved in Fe_2_O_3_-induced inflammatory effect, we further treated THP-1 cells with cytochalasin D, a cytoskeletal inhibitor of endocytosis, before exposure to the Fe_2_O_3_ particles. All cells treated with cytochalasin D showed a decrease in IL-1β release, suggesting that cellular uptake is essential in generating the inflammatory effect (Figure S6). Confocal microscopy was used to study the cellular uptake of fluorescein isothiocyanate (FITC)-labeled Fe_2_O_3_ nanoparticles, and we found that most of the labeled nanoparticles co-localized with an Alexa fluor 594-labeled LAMP1-positive compartment with co-localization coefficients ranging from 73-93% by Image J analysis (Figure 4B). This suggests that Fe_2_O_3_ nanoparticles were mainly taken into the lysosomes of THP-1 cells. TEM data confirmed that Fe_2_O_3_ nanoparticles are encapsulated into vesicular THP-1 compartments of THP-1 cells. Since lysosome is an acidifying environment, Fe_2_O_3_ particles tend to aggregate in this intracellular compartment and interact with its membranes. This likely has led to the lysosome membrane damage due to the reactive surface (nanoplates) or geometric shape of Fe_2_O_3_ nanorods.

In order to understand the biological impact of Fe_2_O_3_ nanoparticles in lysosomal compartments, we asked whether that would impact lysosomal function. Confocal microscopy was used to study the subcellular localization of cathepsin B, a lysosomal enzyme capable of cleaving a Magic Red™-labeled substrate. As shown in Figure S7A, untreated cells show a punctate distribution of Magic Red™, indicating that the enzyme is contained in intact lysosomes. However, after lysosomal damage by MSU, there is a diffuse cytosolic release of the fluorescence marker. Similarly, P3 and R4 nanoparticles induce cathepsin B release, while P1 and R1 nanoparticles are not associated with lysosomal damage. Since cathepsin B is known to contribute to the activation of the NLRP3 inflammasome and IL-1β production^34^, this may explain the severe inflammatory cytokine release in Fe_2_O_3_-treated THP-1 cells. The role of cathepsin B in NLRP3 inflammasome activation was further confirmed by using a cathepsin B inhibitor, CA-074-Me, which shows the inhibitory effect in IL-1β production (Figure S7B). Moreover, we confirmed that active assembly of the NLRP3 inflammasome subunits is required for IL-1β production by using NLRP3- and ASC-gene knockdowns to show the interference in cytokine release in THP-1 cells (Figure 4C).

Based on the mechanism study, we for the first time deciphered the inflammatory pathway of Fe_2_O_3_ in THP-1 cells. As shown in Figure 4D, Fe_2_O_3_ nanoparticles are internalized into lysosomes through endocytosis. Macrophage uptake and lysosomal processing of Fe_2_O_3_ nanoparticles further lead to the interaction with lysosome membrane. Because of the surface reactivity and geometric shape of Fe_2_O_3_ nanoplates and nanorods, these particles may induce lysosome damage, cathepsin B release into cytoplasm, recruitment of NLRP3, pro-caspase 1 and ASC subunits, NLRP3 inflammasome activation and IL-1β release from the macrophages. IL-1β may further participate in a progressive march of inflammation events in organs.

### Examining the effects of Fe_2_O_3_ nanoparticles in cell migration

The nano-SAR profile also indicates that surface reactivity may be responsible for Fe_2_O_3_-induced cell migration. Since monocyte chemoattractant protein-1 (MCP-1 or CC-chemokine ligand 2) is widely reported to be a critical factor for mediating arrest of the monocytic cells and directional migration^35^, we first examined the effects of Fe_2_O_3_ particles on MCP-1 production. After 24 h exposure to Fe_2_O_3_ nanoparticles, significant MCP-1 production was detected in the supernatants of THP-1 cells, and Fe_2_O_3_ plates show higher MCP-1 production than rods (Figure 5A). While P3 stimulates the highest MCP-1 secretion in THP-1 cells, P2 and P4 had moderate effects. Bindarit, as an inhibitor of MCP-1 pathway, could effectively block all Fe_2_O_3_ induced MCP-1 productions. Then we transferred the supernatants of Fe_2_O_3_-treated THP-1 cells to the lower chambers of transwell systems to examine the effects in cell migration. As shown in Figure 5B, Fe_2_O_3_ plates with high surface reactivity induce significant cell migration, and P3 shows the highest level. To investigate whether the macrophage recruitment is a result of MCP-1 production, we examined the effects of supernatants from THP-1 cells exposed to bindarit and Fe_2_O_3_ particles on cell migration. Bindarit treatment results in total reduction of cell migration. These results indicate that Fe_2_O_3_ plates could induce MCP-1 dependent cell migration, and surface reactivity is the dominant property for this effect. Immune cell recruitment is an early statement in acute immune responses and involves transendothelial migration toward the stimulation site to protect health tissues^36^. Thus, with regard to monocyte or leukocyte migration, Fe_2_O_3_ nanoplates display immunostimulatory functions and may serve as a modulator to activate immune cell recruitment.

**Figure 5.**
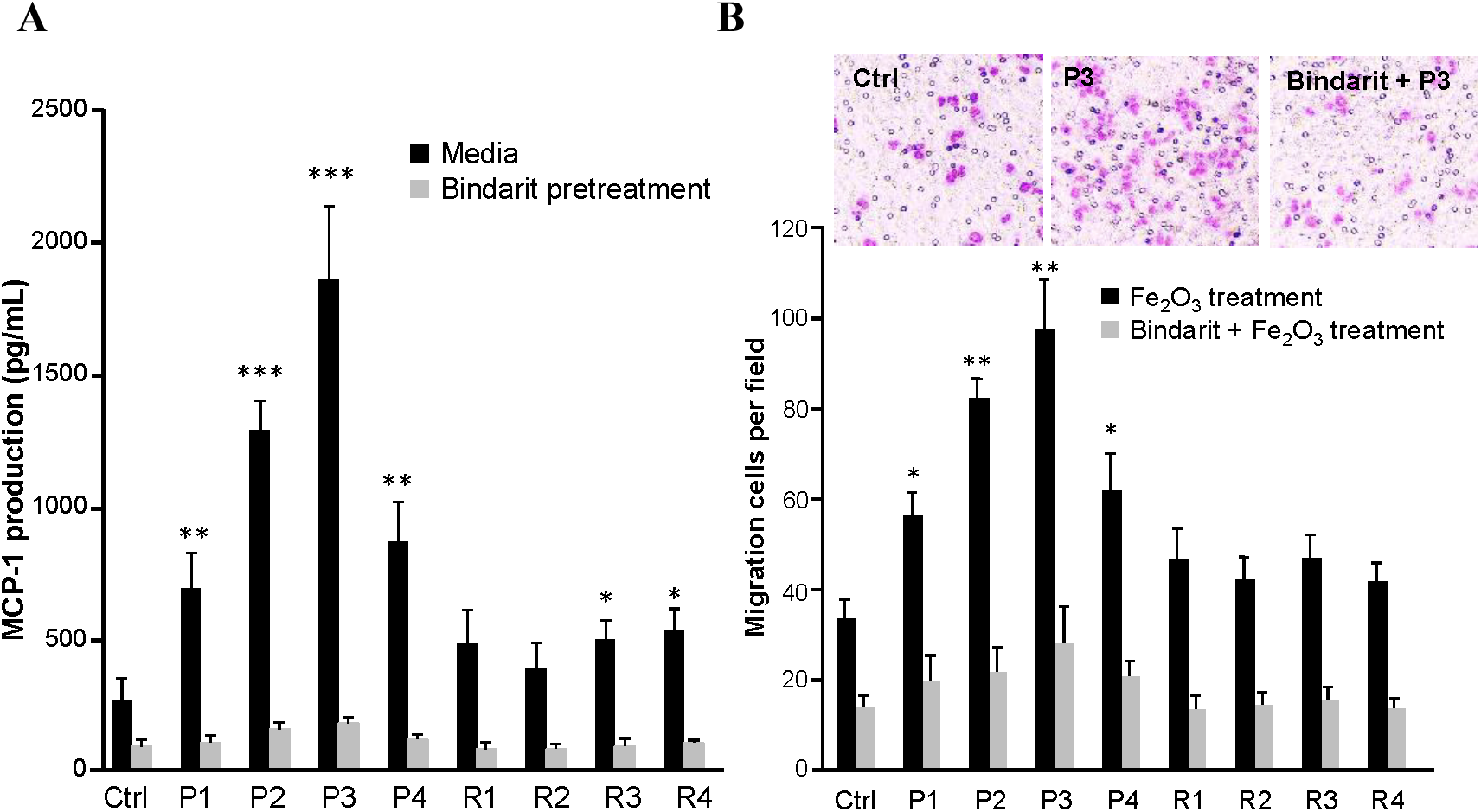
Effects of Fe_2_O_3_ nanoparticles on THP-1 cell migration. **A)** MCP-1 production in the supernatants of Fe_2_O_3_-treated cells, and **B)** transwell migration assays of THP-1 cells. THP-1 cells were pretreated with or without 100 μM bindarit for 3 h before exposure to 100 μg/mL Fe_2_O_3_ nanoparticles. After 24 h incubation, the supernatants of treated THP-1 cells were collected and divided into 2 portions. One was used for MCP-1 measurement by ELISA, and the other one was added into the lower chambers of transwell systems to incubate with THP-1 cells in upper chamber for 24 h. The migration cells were stained with Diff Quick agent and counted under microscope (n=5 images for each treatment). *p< 0.05, **p< 0.01, ***p< 0.001 compared to ctrl (two-tailed Student’s t-test).

### Validation of the inflammatory effects of Fe_2_O_3_ nanoparticles in mouse lungs

In order to further testify the nano-SAR of cellular inflammatory effects induced by Fe_2_O_3_ particles, we used an acute lung injury model to study the effect of oropharyngeal instilled nanoparticles in the whole lung. This study was performed with a particle dose of 2 mg/kg, which has been previously demonstrated to fall on the linear part of the dose-response curve for pulmonary exposure to metal oxide nanoparticles^26, 37^. After 40 h exposure, the animals were sacrificed to collect bronchoalveolar lavage fluid (BALF) and lung tissues. The cytokine release in BALF was determined by ELISA. As shown in Figure 6A, most of the immune cells induced by Fe_2_O_3_ are neutrophils, which are dramatically boosted in P3 and R4 treated animal lungs. In addition, P3 and R4 induced significant cytokine release including IL-1β, TNF-α and LIX (Figure 6B), which is consistent with their *in vitro* inflammatory responses. The migration effect of Fe_2_O_3_ particles was also validated by the MCP-1 production in BALF as well as H&E staining of lung tissues. As shown in Figure 6C, while P3 significantly elevates MCP-1 production and induces massive immune cell recruitment, P1 and R1 have a little effect. Interestingly, R4 exhibits limited effect in MCP-1 production but substantial immune cell recruitment in animal lungs, suggesting there may be other mechanisms involved in animal lungs. All these animal results demonstrated that the nano-SAR in Fe_2_O_3_-induced inflammatory and migration effects could be validated *in vivo*.

**Figure 6.**
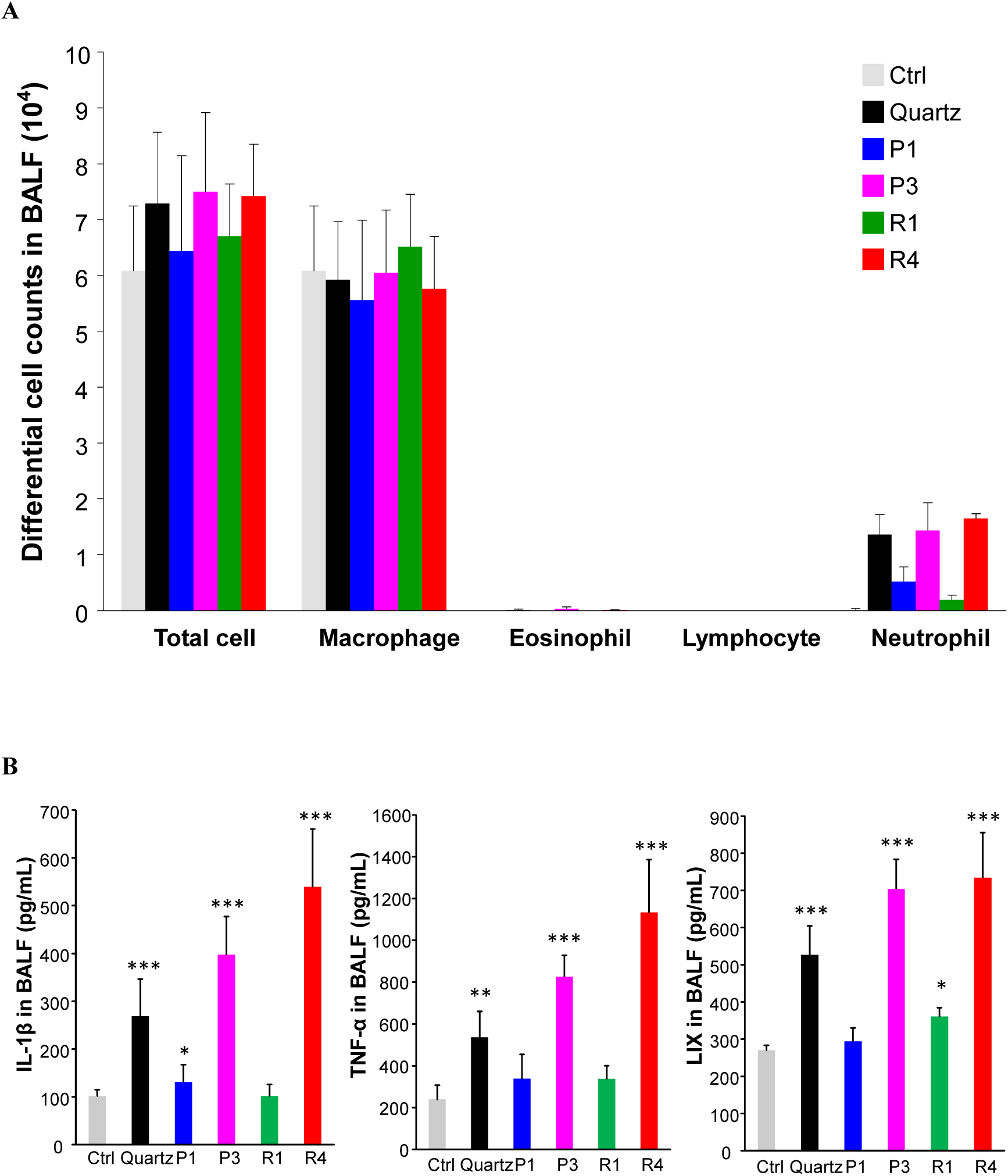
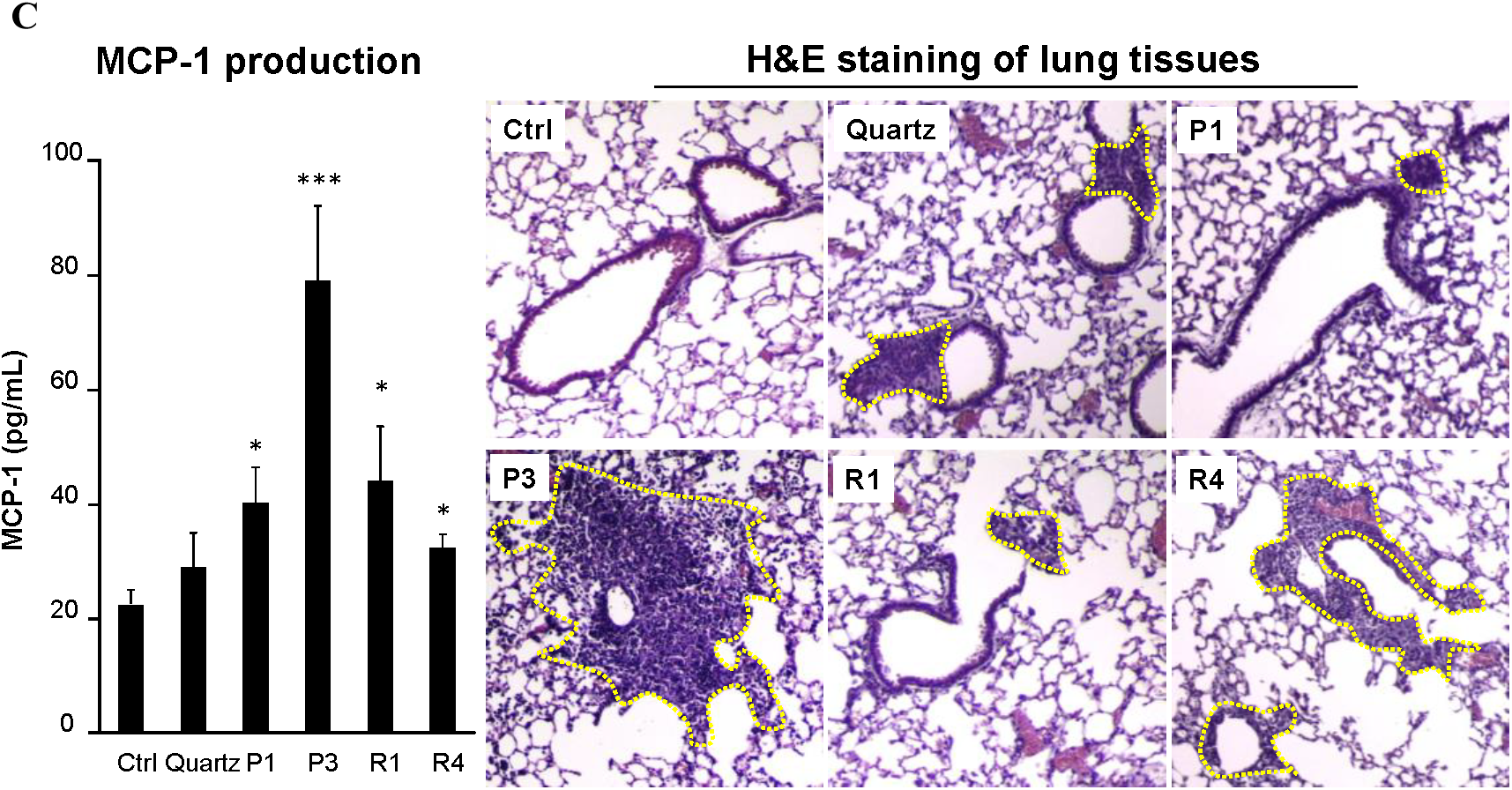
Assessment of inflammatory effects and cell migration in Fe_2_O_3_ treated animal lungs. **A)** Differential cell counts, **B)** cytokine release in BALF, and **C)** MCP-1 production and H&E staining of lung sections from Fe_2_O_3_-treated mice. Selected Fe_2_O_3_ nanoplates (P1, P3) and nanorods (R1, R4) were oropharyngeally administrated at 2 mg/kg (n = 6 mice in each group), while animals received 5 mg/kg quartz exposure were used as positive control. After 40 h, animals were sacrificed to collect BALF for differential cell counting as well as cytokine measurement, including IL-1β, LIX, TNF-α and MCP-1. Data are shown as mean ± s.d. from 4 independent replicates. * p < 0.05,** p < 0.01, *** p< 0.001 compared to vehicle control (two-tailed Student’s t-test). The dashed lines in H&E staining images show the immune cell recruitments.

## Discussions

In this study, we pioneered a multi-hierarchical nano-SAR assessment by simultaneously examining ENM-induced metabolite and protein changes in cells. Unlike traditional method that could only explore the nano-SAR between a specific physicochemical property of ENMs and individual bio-effects^2-4^, a 3D heatmap was established in this study to assess the contributions of 7 physicochemical properties of Fe_2_O_3_ particles to their 6 bio-effects in cells. The nano-SAR investigation could well facilitate the identification of the key physicochemical properties that are responsible for their bio-effects because the biological responses of ENMs have been demonstrated to result from their unique physicochemical properties^3, 38^.

Recently, Zanganeh *et al.* have demonstrated that iron oxide nanoparticle induced pro-inflammatory macrophage polarization plays an important role in tumor therapy^20^, however, the nano-SAR information as well as the detailed inflammatory mechanism shrouds in mystery. We answered these questions by setting up a combinatorial Fe_2_O_3_ library with precisely controlled size and shape as well as systematically assessment of their bio-effects. By regression analysis of particle properties, metabolite and protein changes, aspect ratio and surface reactivity were identified as the key physicochemical properties responsible for the inflammatory effects of Fe_2_O_3_ nanorods and nanoplates, respectively. Fe_2_O_3_-induced cell migration is determined by surface reactivity. These findings were successfully validated in THP-1 cells and animal lungs. *In vitro* experiments further deciphered the MCP-1 dependent cell migration mechanism as well as the NLRP3 inflammatory pathway in Fe_2_O_3_-treated THP-1 cells.

Our study successfully achieved tiered view of the nano-SARs of Fe_2_O_3_ particles in THP-1 cells, providing substantial new insights for tailored design of ENMs by modifying their physicochemical properties to acquire desired bio-effects. This allows us to reduce the toxicities of some hazardous ENMs^7, 8, 28^, or enhance the effects of nanomedicine^5, 6, 39^. Besides of Fe_2_O_3_, the multi-hierarchical nano-SAR assessment method could be potentially extended to other ENMs. This study has far-reaching implications for the sustainable development of nanotechnology.

## Methods

### Materials

Magic Red™ Cathepsin B Assay Kit was purchased from Immunochemistry (Bloomington, MN, USA); Hoechst 33342, H_2_-DCFDA, Alexa fluor 594-labeled goat anti-mouse IgG, anti-LAMP1 primary antibody were purchased from Life Technologies (Grand Island, NY, USA); ELISA kits for detection of murine or human IL-1β, TNF-α, MCP-1 and LIX were purchased from BD biosciences (San Jose, CA, USA); MTS assay kit was purchased from Promega (Madison, WI, USA); bindarit was purchased from Abcam (Cambridge, MA, USA); other chemicals unless stated were purchased from Sigma-Aldrich (St. Louis, MO, USA).

### In-house synthesis of Fe_2_O_3_ nanoparticles

Both Fe_2_O_3_ nanorods and nanoplates were prepared using hydrothermal synthesis methods modified from published procedures^40, 41^. In a typical Fe_2_O_3_ nanorod synthesis, a 10∼16 mL of 0.86 M FeCl_3_•6H_2_O aqueous solution was prepared in a high density polyethylene bottle, to which 4∼10 mL of 1,2-propanediamine solution was added to form a 20 mL synthesis mixture. After stirring for 15 min, the synthesis mixture was transferred to a 23 mL Teflon-lined stainless steel autoclave. The reaction was carried out in an electric oven at 180 °C under autogenous pressure and static conditions. After the reaction was complete, the autoclave was immediately cooled down in a water bath. The fresh precipitate was separated by centrifugation and washed with deionized water and ethanol alternatively for three cycles to remove ionic remnants. The final product was dried at 60 °C overnight under ambient environment. In a typical Fe_2_O_3_ nanoplate synthesis, 0.4325 g of FeCl_3_•6H_2_O was dissolved in 16 mL of ethanol with a small amount of water (1.1-4.0 mL) under vigorous stirring. After FeCl_3_•6H_2_O was completely dissolved, 1.3125 g of sodium acetate was then added and the resulting mixture was mixed for another 15 mins. The reaction and final product collection were carried out the same way as those for the Fe_2_O_3_ naonorods.

### Preparation of Fe_2_O_3_ nanoparticles suspensions in media

The Fe_2_O_3_ stock solutions were prepared by suspending particle powders in DI H_2_O (5 mg/mL) and dispersed in a bath sonicator (Branson, Danbury, CT, USA, model 2510; 100 W output power; 42 kHz frequencey) for 15 min. To prepare the desired concentrations of Fe_2_O_3_ suspensions, an appropriate amount of each Fe_2_O_3_ nanoparticle stock solution was added to cell culture media or PBS, and further dispersed using a sonication probe (Sonics & Materials, USA) at 32 W for 10 s before exposure to cultured cells or animals^42^.

### Physicochemical characterization of Fe_2_O_3_ nanoparticles

A particle suspension (50 μg/mL in DI H_2_O) drop was added on the TEM grids for air-dry at room temperature. The TEM observation was performed on a JEOL 1200 EX instrument (accelerating voltage 80 kV). A Philips X’Pert Pro diffractometer equipped with CuKr radiation were used to obtain the XRD spectra. The hydrodynamic diameter and surface charge in water and cell culture media were measured by dynamic light scattering coupled with zeta potential analyzer (Brookhaven Instruments Corporation, Holtsville, NY, USA). DCF assay was used to evaluate the surface reactivity of Fe_2_O_3_ particles. In detail, 50 μg of H_2_DCF-DA were mixed with 280 μL 0.01 M NaOH and incubated for 30 min at room temperature. The resulting solution was diluted with 1720 μL of a sodium phosphate buffer (25 mmol/L, pH = 7.4) to form 25 μg/mL DCF working solution. A 5 μL aliquots of nanoparticle suspension (5 mg/mL) were added into each well of a 96 multiwell black plate (Costar, Corning, NY), and then 95 μL amount of DCF working solution was added to each well, followed by 2 h incubation. DCF fluorescence emission spectra were recorded by a SpectraMax M5 microplate reader at an excitation wavelength of 490 nm^8^.

### Cell culture and treatment

THP-1 cells and BEAS-2B cells were purchased from ATCC (Manassas, VA, USA), and cultured in RPMI 1640 supplemented with 10% fetal bovine serum and BEGM media, respectively. Before exposure to Fe_2_O_3_ nanoparticles, THP-1 cells were primed by 1 μg/mL phorbol 12-myristate acetate overnight^7^. For cellular exposure, Fe_2_O_3_ nanoparticles were dispersed in complete RPMI 1640 medium at desired concentrations. Control sample was prepared by replacing the nanoparticle suspensions with pure water.

### Nontargeted metabolomics *via* LC-MS

THP-1 cells were exposed to 100 μg/mL Fe_2_O_3_ nanoparticles for 24 h, and 9 cell samples including control and particle treatments were harvested and lyzed in cold lysis buffer (1.5 mL) by a probe sonication. After centrifugation at 20, 000 g for 10 min, the lysis supernatants were collected and extracted by 80% methanol containing 0.5mM L-Methionine-(methyl-13C,d3) and 1mM D-Glucose-1,2,3,4,5,6,6-d7 as internal standards. Dried metabolite pellets were re-suspended in 120 μL of buffer A (0.1% formic acid in 95:5 water/ACN) and 5 μL aliquots were injected for nontargeted LC-MS on a Shimadzu UFLC-XR system (Shimadzu Corporation, Japan) and an AB SCIEX TripleTOF 5600+ system (AB SCIEX, Foster City, CA). Samples were separated on a C18 reversed phase HPLC column (2.1mm × 100mm, 100Å, 1.7μm, Waters, Milford, MA) at 350 μL/min with a liner gradient of buffer A and buffer B (100% ACN) as follows: isocratic conditions at 100% A (0%B) for 1min, a linear gradient from 100%A (0%B) to 30% A (70%B) over 2 min, a linear gradient from 30%A (70%B) to 0% A (100%B) over 7 min,isocratic conditions at 100% B for 2 min. The column temperature maintained at 50°C.

To acquire the MS data, the mass spectrometry conditions were set as follows: the ion spray voltage was +5.5kV (positive ion mode) or −4.5kV (negative ion mode); turbo spray temperature was 550 °C; nebulizer, heater and curtain gases were at 50, 50 and 30 psi, respectively; TOF MS was scanned at the mass range of m/z 50 ∼ 1200. Analyst v1.6.0 software (AB Sciex) was used to collected raw data, which was further converted into mzXML data format by proteoWizard software (Spielberg Family Center for Applied Proteomics, Los Angeles, CA) for further data processing.

The XCMS platform (https://metlin.scripps.edu/xcms/) was used for peak detection, retention time collection and alignment. All data-collection parameters were set to the “UPLC Triple TOF” default values. Retention times (RT), m/z values and peak intensities of metabolites were exported to an Excel spreadsheet for processing. The peak intensities were normalized to the internal standards: L-Methionine-(methyl-13C, d3) (m/z, 154.077) for positive mode and D-Glucose-1,2,3,4,5,6,6-d7 (m/z, 186.099) for negative mode. Preprocessed data sets were analyzed using Matlab (MathWorks, Natick, MA) and Metaboanalyst (www.metaboanalyst.ca) to perform scatter plot, heat map, cluster and ANOVA analysis. Euclidean distance (ED) is used to measure the dissimilarity of samples with multivariate variables^29^. Here, the ED between control and nanoparticle-treated samples is defined by the length between the two cluster centers. It was calculated by SPSS using following formula: 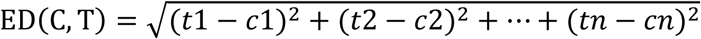, where tn and cn are the log-transformed normalized peak intensities of metabolite n in nanoparticle-treated sample and control sample, respectively (n=2854).

### Proteomics *via* nanoflow LC-MS/MS

The cell samples exposed to Fe_2_O_3_ nanoparticles were harvested and lyzed. After centrifugation at 20000 g, the lysis supernatants (1 mL) were added to 8 mL extraction solute (acetone/ethanol/acetic acid 50:50:0.1), which was pre-cooled at −20 °C. The mixture was stored at 4 °C for 24 h. Then the proteins were collected by centrifugation at 20000 g for 30 min. After re-dissolution, the protein concentrations in all the samples were equivalent by a Bradford method, and then the protein samples were denaturalized, digested and desalted according to a reported method^44^. Finally, the resulting peptide samples were re-dissolved in 200 μL 0.1% formic acid for nano-LC-MS/MS analysis.

A LTQ Orbitrap Velos was equipped with an Accela 600 HPLC system (Thermo, San Jose, CA) to establish the nano-LC-MS/MS system. The peptide samples were injected into a capillary trap column (200 mm i.d. × 4 cm, 120 Å, 5 mm), and then separated on an analytical column (15 cm × 75 mm i.d., 3 mm, 120 Å). Both of the columns were packed with C18 AQ beads. The separation buffer consisted of 0.1% (v/v) formic acid in DI H_2_O (buffer A) and 0.1% (v/v) formic acid in acetonitrile (ACN) as buffer B. The nano-LC-MS/MS analysis was performed based on a previously reported method^43^. All of the mass spectra were collected in a data dependent mode.

The resulted raw files were searched in MaxQuant (Version1.3.0.5) using Integrated Uniprot protein fasta database of human. Peptide searching was constrained using fully tryptic cleavage, allowing less than 2 missed cleavages sites for tryptic digestion. Variable modifications included methionine oxidation, acetylation of protein N-term and phosphorylation (STY). Precursor ion and fragment ion mass tolerances were set as 5 ppm and 0.08 Da, respectively. The false discovery rate (FDR) for peptide and protein were less than 1% and peptide identification required a minimum length of six amino acids. The cell pathways and functions related to the identified differential proteins were explored by KEGG (http://www.kegg.jp/) and UniprotKB (http://www.uniprot.org/help/uniprotkb) database.

### IL-1β and TNF-alpha Detection by ELISA

IL-1β and TNF-alpha productions were detected in the culture media of THP-1 cells using human IL-1β and TNF-alpha ELISA Kit (BD; San Jose, CA, USA). Briefly, aliquots of 5×10^4^ THP-1 cells were seeded in 0.1 mL complete medium and primed with 1 μg/mL phorbol 12-myristate acetate (PMA) overnight in 96-well plates (Corning; Corning, NY, USA). Cells were treated with the desired concentration of the particle suspensions made up in complete RPMI 1640 medium, supplemented with 10% fetal bovine serum and 10 ng/mL lipopolysaccharide (LPS).

### Cell migration assay

THP-1 cells were pretreated or not with 100 μM bindarit for 3 h before exposure to 100 μg/mL Fe_2_O_3_ nanoparticles. After 24 h incubation, the supernatants were collected for MCP-1 detection or cell migration assay, which was performed in 24-transwell plates with polycarbonate membranes of 8 μm pores (Corning, NY, USA). Lower wells were filled with 500 μL aliquots of the collected supernatants, and 2 x10^5^ THP-1 cells (100μL) were seeded into each of the upper wells. After incubation for 6h, nonmigrated cells were scraped off from the upper side of the membrane and cells remaining within the pores or below the membranes were stained with Diff Quick^35^. Cell numbers were calculated under microscope by randomly selecting at least 5 individual fields for each sample.

### TEM imaging of Fe_2_O_3_ particles in THP-1 cells

After exposure to 25 μg/mL Fe_2_O_3_ for 24 h, THP-1 cells were collected, washed and fixed with 2% glutaraldehyde in PBS. After 1 h post-fixation staining in 1% osmium tetroxide, a dehydration process was performed by treating the cells in a graded series of ethanol, propylene oxide, and finally the cell pellets were embedded in Epon. A Reichert-Jung Ultracut E ultramicrotome was used to cut the TEM sections with approximately 50-70 nm thickness. The sections were further stained with uranyl acetate and Reynolds lead citrate before examining on TEM as previously reported^42^.

### Confocal microscopy imaging

Leica confocal SP2 1P/FCS microscopes were used to visualize Fe_2_O_3_ uptake and cathepsin B release in THP-1 cells. High magnification images were obtained under the 63X objective. To visualize the cellular distribution, THP-1 cells were treated with 25 μg/mL FITC-labeled Fe_2_O_3_ nanoparticles for 6 h, fixed and stained with Hoechst 33342 and Alexa Fluor 594 labeled antibodies to visualize nuclei and lysosomes, respectively. For cathepsin B imaging, cells exposed to 100 μg/mL Fe_2_O_3_ particles for 16 h were stained with Magic Red™ Cathepsin B kit and Hoechst 33342 for confocal microscopy imaging.

### Inflammation test in mouse lungs

Mice were exposed to nanoparticle suspensions using oropharyngeal aspiration at 2 mg/kg. Eight-week-old male C57Bl/6 mice purchased from Soochow University were used for animal experiments. All animals were housed under standard laboratory conditions that have been set up according to Soochow University guidelines for care and treatment of laboratory animals. These conditions were approved by the Chancellor’s Animal Research Committee at Soochow University and include standard operating procedures for animal housing (filter-topped cages; room temperature at 23 ± 2 °C; 60% relative humidity; 12 h light, 12 h dark cycle) and hygiene status (autoclaved food and acidified water). Animals were exposed by oropharyngeal aspiration as described by us^8^. Briefly, animals were anesthetized by intraperitoneal injection of ketamine (100 mg/kg)/xylazine (10 mg/kg) in a total volume of 100 μL. The anesthetized animals were held in a vertical position. 50 μL aliquots of the nanoparticle suspensions in PBS were instilled at the back of the tongue to allow pulmonary aspiration of a dose of 2 mg/kg. Each experiment included control animals, which received the same volume of PBS. The positive control in each experiment received 5 mg/kg quartz. Each group included six mice. The mice were sacrificed after 40 h exposure. BALF and lung tissue were collected as previously described. The BALF was used for performance of total and differential cell counts and measurement of IL-1β, TNF-α, MCP-1 and LIX levels. Lung tissue was stained with hematoxylin/eosin.

### Statistical Analysis

All the experiments were repeated at least thrice with 3-6 replicates. Error bars represent the standard deviations (s.d.). All the cell and animal samples were randomly allocated into experimental groups by drawing lots. All the experiments were repeated at least thrice with 3–6 replicates. Results were expressed as mean ± SD of multiple determinations from at least three separate experiments. One-way ANOVA or Student t test was used for statistical analysis in MetaboAnalyst 3.0 and excel 2010. The difference is regarded statistically significant with p ≤0.01 and FDR≤0.05. Correlation analysis of the differential proteins and metabolites was performed in Matlab R2009b. The correlation is considered as statistically significant with correlation coefficient (r) ≤-0.8 or r≥0.8, and p ≤ 0.01. The Euclidean distance (ED) and linear regression analysis of the EDs and particle properties were achieved in SPSS 18.0. The 3D profile of the Structure-Activity relationships was also done in Matlab.

### Data availability

The data that support the plots within this paper and other findings of this study are available from the corresponding author on reasonable request.

## Acknowledgements

This work was supported by the grant from the National Natural Science Foundation of China (No. 31671032), Key Project of Natural Science Foundation of the Higher Education Institutions of Jiangsu Province (No. 17KJA310003), and a project funded by the Priority Academic Program Development of Jiangsu Higher Education Institutions (PAPD). R.L. is supported by the recruitment program of Global Youth Experts of China.

## Author contributions

X.C., Z.J. and R.L. conceived and designed the study; X.C. did most experiments; Z.J. synthesized the Fe_2_O_3_ nanoparticles, and performed TEM, XRD, Zeta potential and hydrodynamic size characterization with H.Z.; J. D. performed the LC-MS analysis for metabolomics study; F. W. and J. L. contributed to the proteomics analysis. The writing of the paper was led by X. C. and R.L with participation from Z.J. and C.K.

## Additional information

Supplementary information is available in the online version of the paper. Reprints and permissions information is available online at www.nature.com/reprints. Publisher’s note: Springer Nature remains neutral with regard to jurisdictional claims in published maps and institutional affiliations. Correspondence and requests for materials should be addressed to R.L. and Z.J.

